# Early Intervention with Electrical Stimulation Reduces Neural Damage After Stroke in Non-human Primates

**DOI:** 10.1101/2023.12.18.572235

**Authors:** Jasmine Zhou, Karam Khateeb, Azadeh Yazdan-Shahmorad

## Abstract

Ischemic stroke is a neurological condition that results in significant mortality and long-term disability for adults, creating huge health burdens worldwide. For stroke patients, acute intervention offers the most critical therapeutic opportunity as it can reduce irreversible tissue injury and improve functional outcomes. However, currently available treatments within the acute window are highly limited. Although emerging neuromodulation therapies have been tested for chronic stroke patients, acute stimulation is rarely studied due to the risk of causing adverse effects related to ischemia-induced electrical instability. To address this gap, we combined electrophysiology and histology tools to investigate the effects of acute electrical stimulation on ischemic neural damage in non-human primates. Specifically, we induced photothrombotic lesions in the monkey sensorimotor cortex while collecting electrocorticography (ECoG) signals through a customized neural interface. Gamma activity in ECoG was used as an electrophysiological marker to track the effects of stimulation on neural activation. Meanwhile, histological analysis including Nissl, cFos, and microglial staining was performed to evaluate the tissue response to ischemic injury. Comparing stimulated monkeys to controls, we found that theta-burst stimulation administered directly adjacent to the ischemic infarct at 1 hour post-stroke briefly inhibits peri-infarct neuronal activation as reflected by decreased ECoG gamma power and cFos expression. Meanwhile, lower microglial activation and smaller lesion volumes were observed in animals receiving post-stroke stimulation. Together, these results suggest that acute electrical stimulation can be used safely and effectively as an early stroke intervention to reduce excitotoxicity and inflammation, thus mitigating neural damage and enhancing stroke outcomes.

## Introduction

Ischemic stroke is a major type of brain injury that results in high mortality and serious long-term disability for adults, especially in the aging population. Globally, over 7.6 million people suffer from ischemic stroke each year, causing significant health and economic burdens worldwide (1, 2). An ischemic stroke happens when blood flow within the brain is interrupted, leading to a lack of oxygen supply, energy depletion, and subsequent neuronal death. Acute intervention within hours after stroke onset offers the most critical therapeutic opportunity as it can reduce irreversible tissue injury, resulting in improved neurological and functional outcome for stroke patients (3). However, currently available treatments within the acute window are highly limited, and approved interventions such as the administration of tissue plasminogen activator (t-PA) and catheter-based thrombectomy often have strict patient selection criteria (3, 4). During the past few decades, there has been a large amount of experimental research and clinical trials on neuroprotective drug therapies for acute ischemic stroke, with the aim to interrupt the ischemic cascade and thereby reduce neuronal death (5–7). However, most of the drugs failed to show consistent clinical efficacy when moving from animals to humans (8–10). Therefore, there is a pressing need to expand the therapeutic options for acute ischemic stroke and improve the translation from bench to bedside to help millions of stroke patients retain the maximum quality of life.

In recent years, novel neural modulation paradigms such as electrical brain stimulation have been proposed as a promising treatment for ischemic stroke. Most of these stimulation paradigms target the subacute or chronic phase of stroke to promote neural plasticity and functional recovery, rather than reducing permanent ischemic damage (11–16). As a result, it might take months of treatment in conjunction with rehabilitative training for only a subset of patients to see positive results from these interventions (17–19). Meanwhile, comparing to the chronic implementation of electrical stimulation for stroke, the acute application of such strategy is still relatively rare, especially in larger animal models and humans due to the risk of causing adverse effects and greater tissue damage related to ischemia-induced electrical instability and spreading depolarizations (SDs). It has been widely reported that perilesional tissues are particularly susceptible to SDs, marked by intense neuronal depolarization waves that can lead to increased metabolic stress, neuronal swelling, and lesion progression (20–23). However, despite the concern about SDs, a few studies have successfully demonstrated the use of sensory or direct brain stimulation to exert neuroprotection and reduce tissue damage for rodents with acute stroke (24–28). Similar to neuroprotective drugs, these acute stimulation strategies aim to reduce the irreversible damage caused by the ischemic cascade within hours after stroke onset or after reperfusion by reducing inflammation, apoptosis, oxidative stress, and excitotoxicity, thus preserving the perilesional tissues surrounding the ischemic core (29, 30). While these results from rodent studies are promising, the scale and anatomical differences between rodent and human brains still hinder the feasibility of clinical translation. We need a more comprehensive understanding of how electrical brain stimulation drives changes in the physiology of neuronal networks at scales comparable to human brains before these strategies can be successfully translated from bench to bedside for stroke patients.

As a result, two major gaps in knowledge need to be addressed as we seek to implement brain stimulation paradigms for acute ischemic stroke: 1) The protective effect of electrical stimulation needs to be evaluated across large cortical areas in a more clinically relevant animal model. 2) We need to have a better understanding of the mechanisms underlying stimulation-induced changes, from both electrophysiological and cellular perspectives. In this study, we used a novel set of approaches capable of addressing these two gaps to investigate stimulation-induced neuroprotection. We combined a lesion-based toolbox (31, 32), state-of-the-art neurophysiology techniques, and a range of histology markers to study the neuroprotective effects of cortical electrical stimulation following acute ischemic stroke in non-human primates (NHPs). We compared multiple aspects of physiological responses to stimulation from large areas (∼3 cm^2^) of the macaque sensorimotor cortex at up to 4 hours after stroke onset to interrogate the mechanisms underlying any observed neuroprotection. The unprecedented insights gained from these experiments will inform the development of next-generation brain stimulation paradigms that can be used as an alternative treatment in the acute window after stroke to minimize neuronal damage, reduce severe disabilities, and improve functional outcomes for stroke patients.

## Results

### Reduction in ischemic lesion volume with acute cortical stimulation

In this study, we induced controlled cortical lesions in the primate sensorimotor cortex using the photothrombotic technique (31–33), which generated focal infarcts by photo-activation of a light-sensitive dye that interrupts local blood flow. The infarct size and location are controlled by setting a constant illumination intensity and aperture size through an opaque light mask placed above the cortical surface (Fig. 1A-B). In control monkeys D and E, we collected ECoG data through our customized multi-modal interface (Fig. 1B), which includes baseline before lesioning, during lesion induction, and around 3 hours post lesioning to monitor the network dynamics around the injury site. In stimulated monkeys F and G, we recorded 1 hour after lesioning and then applied electrical stimulation at approximately 8 mm medial to the lesion center on the ipsilesional hemisphere (Fig. 1B-C, blue arrow). The stimulation trains employed a theta burst paradigm with five 1kHz pulses within each burst. Each stimulation block lasted 10 minutes, with recordings of spontaneous activity in between the blocks to track changes in neurophysiology as stimulation continued (Fig. 1D).

**Figure 1.**
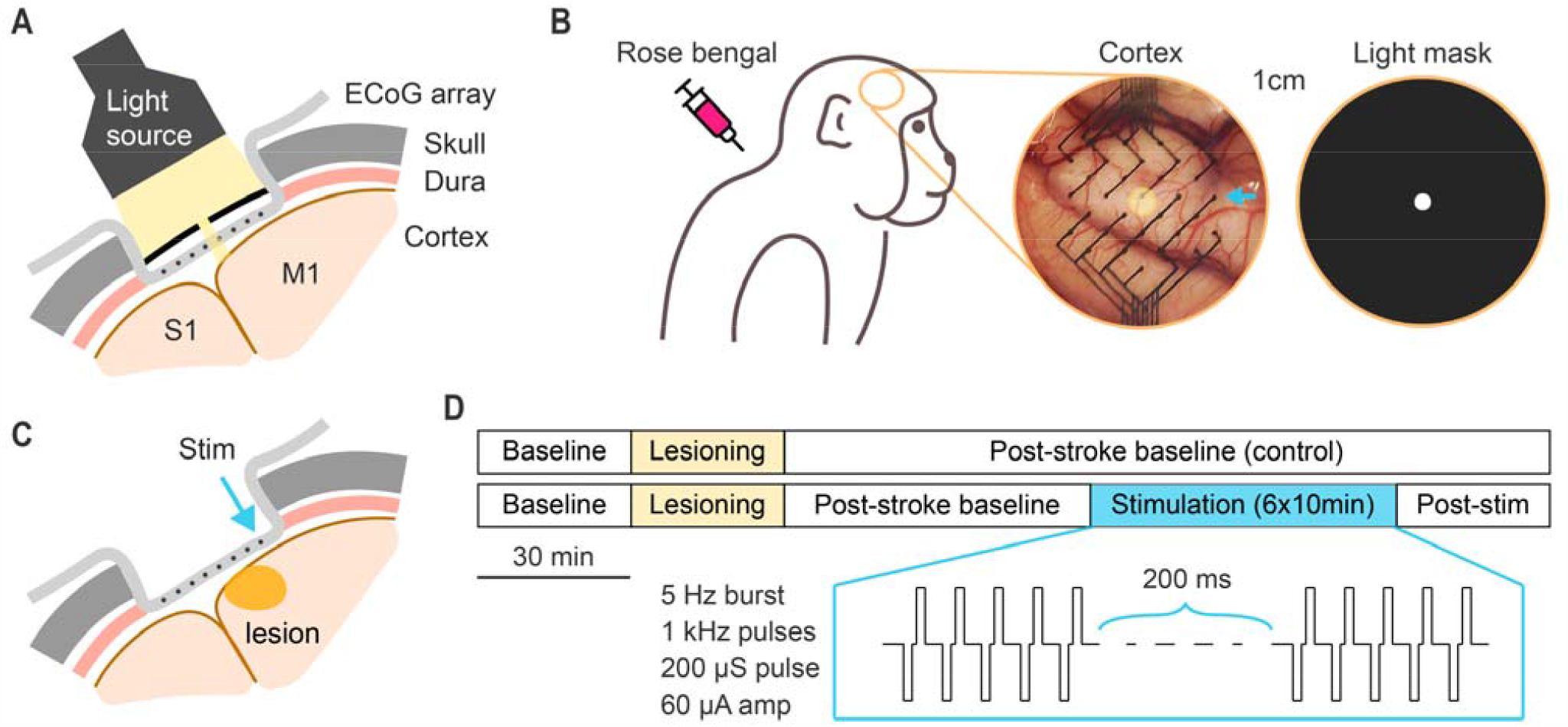
Schematics of experimental procedures. **A**. Illustration of the method to induce photothrombotic lesion in the NHP cortex. **B**. Cortical view of the multi-model artificial dura used to record ECoG. Yellow circle indicates the area illuminated with light source. Blue arrow points to the stimulation electrode. **C**. Illustration of the location of induced lesion and electrical stimulation. **D**. Experimental timeline for the control (monkey D and E) and stimulated groups (monkey F and G).

To investigate the neuroprotective effects of electrical stimulation following acute ischemic stroke, we first quantified the extent of ischemic injury within the sensorimotor cortex through histological analysis. Specifically, we performed Nissl staining on mounted coronal sections of the brain to evaluate the amount of cell death and estimate lesion volumes in both control (monkey D, E) and stimulated animals (monkey F, G). The loss of Nissl substance at the infarct core led to distinct pale areas and well-defined boundaries on the stained sections (Fig. 2A). Using this identified lesion boundary, we applied edge detection and linear interpolation between sections to reconstruct the brain with the ischemic lesion in 3D space (Fig. 2B), including distinct anatomical features of the sensorimotor cortex such as the central sulcus. This reconstruction was used to estimate the lesion volumes in each animal (Fig. 2C). We found that in control monkeys D and E that did not receive stimulation post-stroke, the estimated lesion volumes were 35.3 and 28.4 mm^3^ respectively, while in the stimulated monkeys F and G, the lesion volumes were 20.3 and 15.9 mm^3^ respectively. Noticeably, the ischemic lesions in stimulated animals were smaller than controls in both their depth and medial-lateral width (Fig. 2C-D). As the stimulation pulses were also delivered medially from the infarct core on the medial-lateral axis (Fig. 1C and 2B), these results suggest acute electrical stimulation post stroke may have reduced the infarct size by protecting the brain from ischemic injury progression during the 4 hours after lesioning.

**Figure 2.**
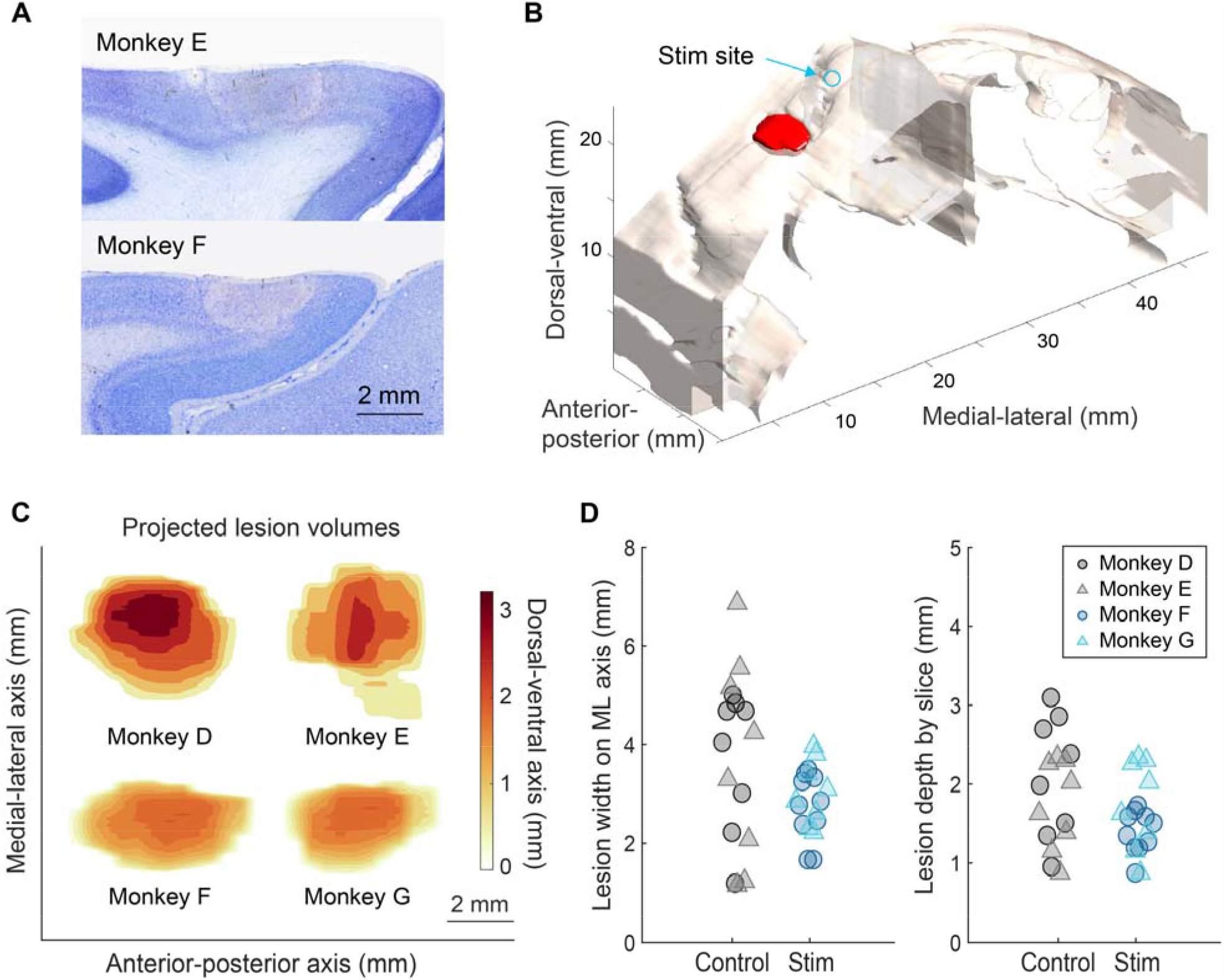
Histology analysis of ischemic damage and lesion size. **A**. Examples of Nissl-stained coronal slices with distinct lesion boundaries. **B**. 3D reconstruction of the cortex and lesion (in red) for monkey F. Blue arrow points to the stimulation electrode. **C**. Estimated lesion volume comparison for monkeys D to G. **D**. Slice by slice comparison of lesion widths and depths for the control (monkeys D and E) and stimulated group (monkeys F and G).

### Decrease of post-stroke neural activity by acute electrical stimulation

To further understand the physiological mechanisms underlying the reduced infarct volume as a result of acute electrical stimulation, we incorporated the histological results with the analysis of electrophysiology recordings by identifying the electrodes that overlapped with the reconstructed lesion . We registered the ECoG electrodes found to be within the anatomically defined lesion by overlaying the surgical image and the reconstructed cortex based on the location of and distance between sulci (Fig. 3A). After classifying the ipsilesional electrodes in each animal as either lesion or non-lesion, we computed the gamma band (30-59 Hz) signal power at each electrode, since gamma rhythms in the brain have been found to be correlated with neural activity levels and firing rate (34, 35). By visualizing the gamma power changes on heatmaps (Figure 3B), we found that in both control and stimulated monkeys, the lesion electrodes showed decreased gamma power as early as 10 minutes after ischemic lesioning, which persisted throughout the experiment (up to 3 hours). This observation confirmed the location of ischemic injury and neuronal death caused by photothrombosis. In addition, we observed a gradual but large-scale downregulation of gamma power across the entire ipsilesional sensorimotor region in response to post-stroke stimulation for monkeys in the stimulation group (Fig. 3B, bottom: showing monkey G as an example). This was distinctively different from what was observed in the control monkeys, as gamma power at some of the non-lesion electrodes was elevated above the baseline at around 90 minutes post lesioning (Fig. 3B, top: showing monkey E as an example), suggesting hyperactivation of perilesional areas as a result of focal ischemia. Similar trends can also be seen from the bar plots grouping two control and two stimulated monkeys (Fig. 3C), in which the gamma power at non-lesion electrodes increased from 50 minutes to 90 minutes post-stroke in monkeys D and E while decreased in monkeys F and G (paired sample t-test with Bonferroni corrections for multiple comparisons, family-wise error rate of 0.05). Together, these results showed that gamma power was mildly suppressed across the injured sensorimotor cortex as a result of continuous electrical stimulation, suggesting that stimulation delivered from 60 minutes after stroke may prevent excessive neuronal activation and energy depletion caused by the ischemic cascade, thus slowing the lesion progression and reducing neuronal death.

**Figure 3.**
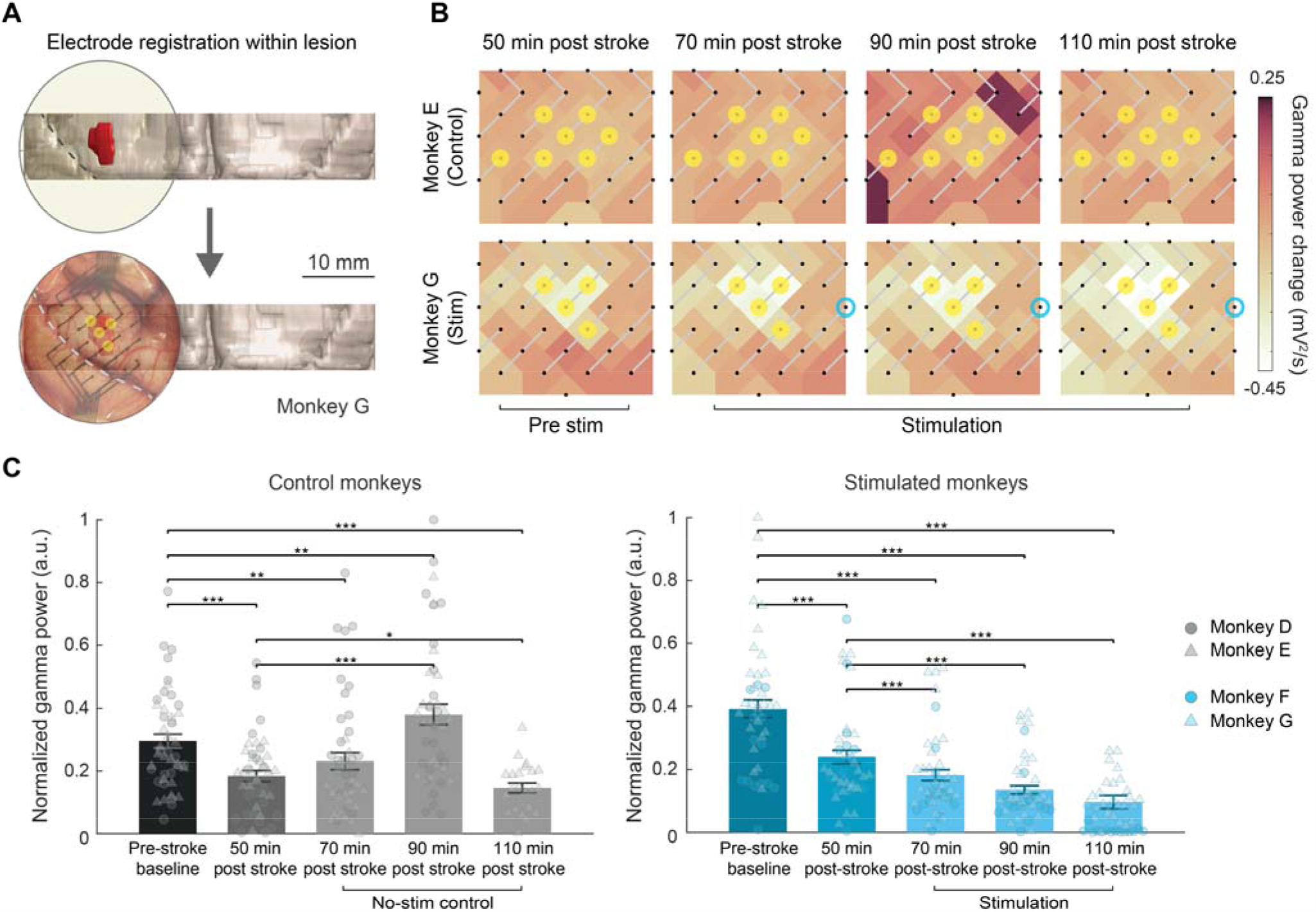
Electrophysiology analysis of the ipsilesional sensorimotor cortex after stroke and electrical stimulation. **A**. Electrode registration within the histologically defined lesion. Yellow circles indicate lesion electrodes. **B**. Heatmaps of the changes in gamma ECoG power for one control (monkey E) and one stimulated animal (monkey G). Yellow circles indicate lesion electrodes. Blue circles indicate stimulation electrode. **C**. Summary and statistics for gamma ECoG power in the control (monkeys D and E) and stimulated group (monkeys F and G) at different time points after stroke and stimulation.

### Decrease of post-ischemia cFos activity by acute electrical stimulation

To investigate the downregulation of neural activity and its potential neuroprotective effects, we went back to histological tools and performed immunohistochemistry staining of c-Fos, a common cellular marker of neuronal activity. c-Fos is a protein derived from the immediate early gene *c-fos*, which gets transiently expressed in neurons quickly following depolarization in the cerebral cortex, and can be detected reliably using immunohistochemistry up to many hours after intense neuronal activation in primates (36, 37). Meanwhile, it was also widely reported that c-Fos expression can be temporarily elevated following focal ischemic stroke as part of the injury response (38, 39). Therefore, we applied immunostaining against c-Fos protein on coronal sections of the ipsilesional sensorimotor cortex. These sections were also co-stained with antibodies against the neuronal nuclear marker NeuN to establish precise lesion boundaries for the subsequent c-Fos analysis. For each control (monkey C, D) and stimulated animal (monkey F, G), we stained two coronal sections around the center of the lesion. On each stained section, two images were taken at the medial and lateral side of the ischemic core respectively, at 0-1 mm and 2-3 mm from the lesion boundary (Fig. 4A). We saw that in control monkeys C and D, there is very high level of c-Fos expression both immediately adjacent to (0-1mm) and a small distance away from (2-3 mm) the lesion boundary (Fig. 4B, top). By contrast, we only saw mild c-Fox expression in stimulated monkeys adjacent to the lesion and extremely low expression at a distance away (Fig. 4B, bottom). After calculating the density of c-Fos positive cells per unit area, we found that cFos activation is significantly higher at a shorter distance from the lesion boundary (Two-way ANOVA, p<0.05). More importantly, the level of c-Fos was much higher in the control monkeys compared to the stimulated monkeys (Two-way ANOVA, p□<□0.001) (Fig. 4C). These results suggest that electrical stimulation applied acutely after stroke reduced the intensity of neuronal depolarization and potentially prevented further cell death caused by overactivation of cortical neurons and excitotoxicity surrounding the ischemic core.

**Figure 4.**
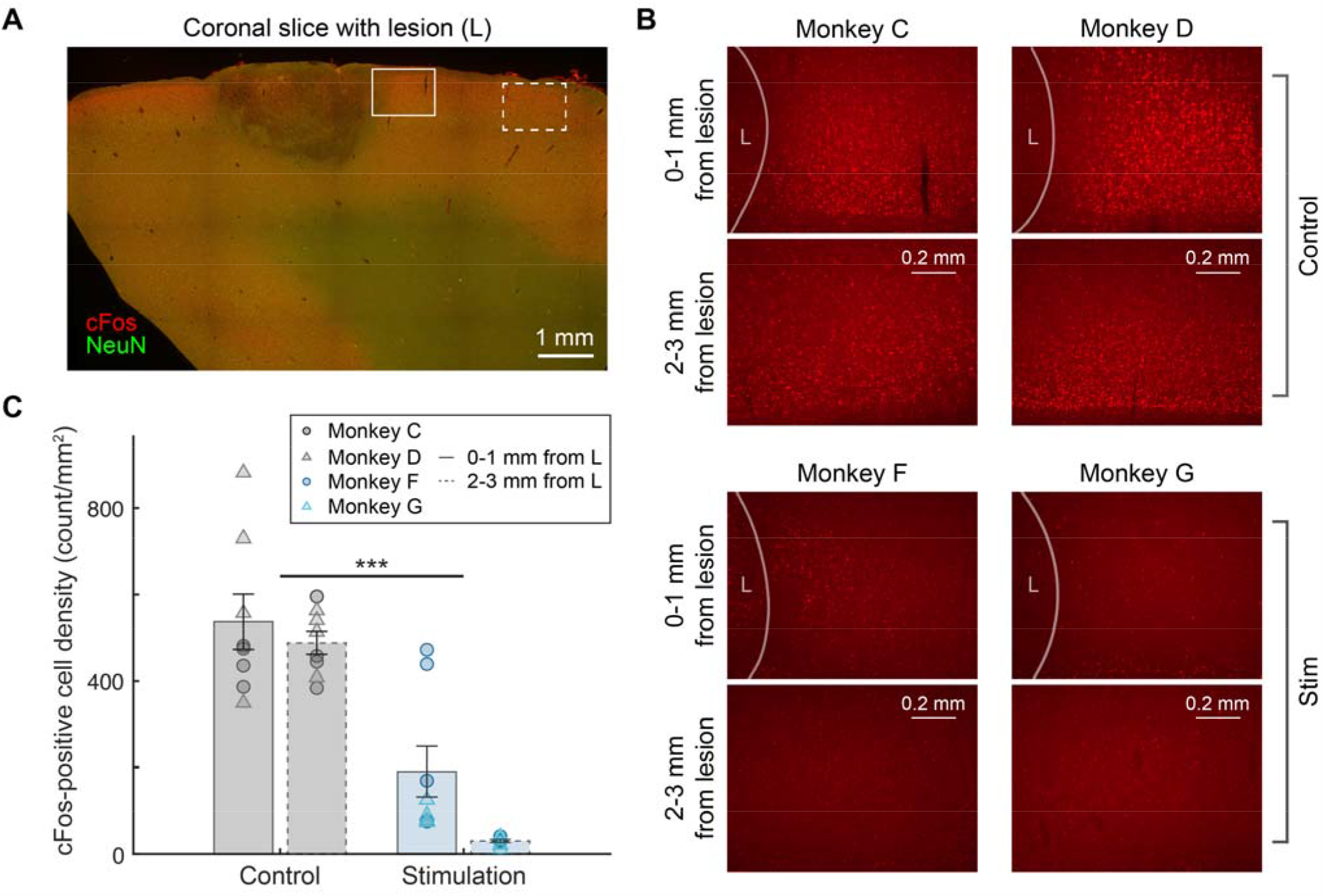
Immunohistochemistry staining of c-Fos protein. **A**. Example coronal section co-stained with cFos (red) and NeuN (green). Solid rectangle represents the area 0-1 mm from lesion boundary and dashed rectangle indicates 2-3 mm from lesion boundary. **B**. Example c-Fos staining from monkeys C, D, F, and G at 0-1 and 2-3 mm from the lesion boundary respectively. **C**. Summary of estimated c-Fos positive cell density for the control (monkeys C and D) and stimulated group (monkeys F and G) at 0-1 and 2-3 mm from the lesion boundary. Two-way ANOVA detected a significant effect of electrical stimulation that leads to reduced c-Fos activation (***: p<0.001).

### Alleviating neuroinflammatory response through acute electrical stimulation

To understand the role of electrical stimulation in reducing excitotoxicity and neuronal death caused by acute ischemic lesion, we evaluated the neuroinflammatory response surrounding the ischemic core by quantifying microglia activation and accumulation in the peri-infarct region. Glial cells including microglia are known to play a significant role in the ischemic cascade and glutamate excitotoxicity (40). Therefore, we applied immunostaining against ionized calcium-binding adapter molecule 1 (Iba1), a widely used intracellular microglial marker (41), on ipsilesional sections of the sensorimotor cortex. Again, these sections were co-stained with antibodies against NeuN to define the lesion boundaries and select region of interests (ROIs) for all subsequent image analysis. For monkeys C, D, F, and G respectively, we stained two coronal sections per animal, and took four ROIs per section at the medial and lateral sides of the ischemic core (Fig. 5A). We observed stronger expression of Iba1 and greater number of Iba1-positive cells in the control animals compared to stimulated animals. Specifically, we saw a greater percentage of microglia with activated morphology in the controls, as characterized by larger cell body area and reduced ramification (Fig. 5B, white arrows). After comparing the binarized ROIs from control and stimulation groups, we found greater level of microglial activation in the controls reflected by greater percentage coverage of Iba1 staining (Fig. 5C. One-way ANOVA and Bonferroni corrections for multiple comparisons, p<0.001). Through cell segmentation, we also found that microglia density near the lesion boundary is higher in the control monkeys C and D than in the stimulated monkeys F and G (Fig. 5D. One-way ANOVA and Bonferroni corrections for multiple comparisons, p<0.001). Note that ipsilesional regions with smaller lesions in control monkey C also showed high level of microglial reactivity reflected by both activation and density, suggesting that the decrease in microglial response was not a direct result of the variation in lesion sizes. Combining these results with the smaller lesion volumes, reduced c-Fos expression, and downregulated neural activity in animals receiving stimulation, our findings suggest that post-stroke electrical stimulation can exert neuroprotection on the sensorimotor cortex by alleviating the inflammatory response, reducing excitotoxicity and conserving critical metabolic energy. Taking advantage of this protective effect, cortical electrical stimulation can be investigated further as a promising therapeutic intervention in the acute phase after stroke to minimize the irreversible neuronal damage caused by the ischemic cascade.

**Figure 5.**
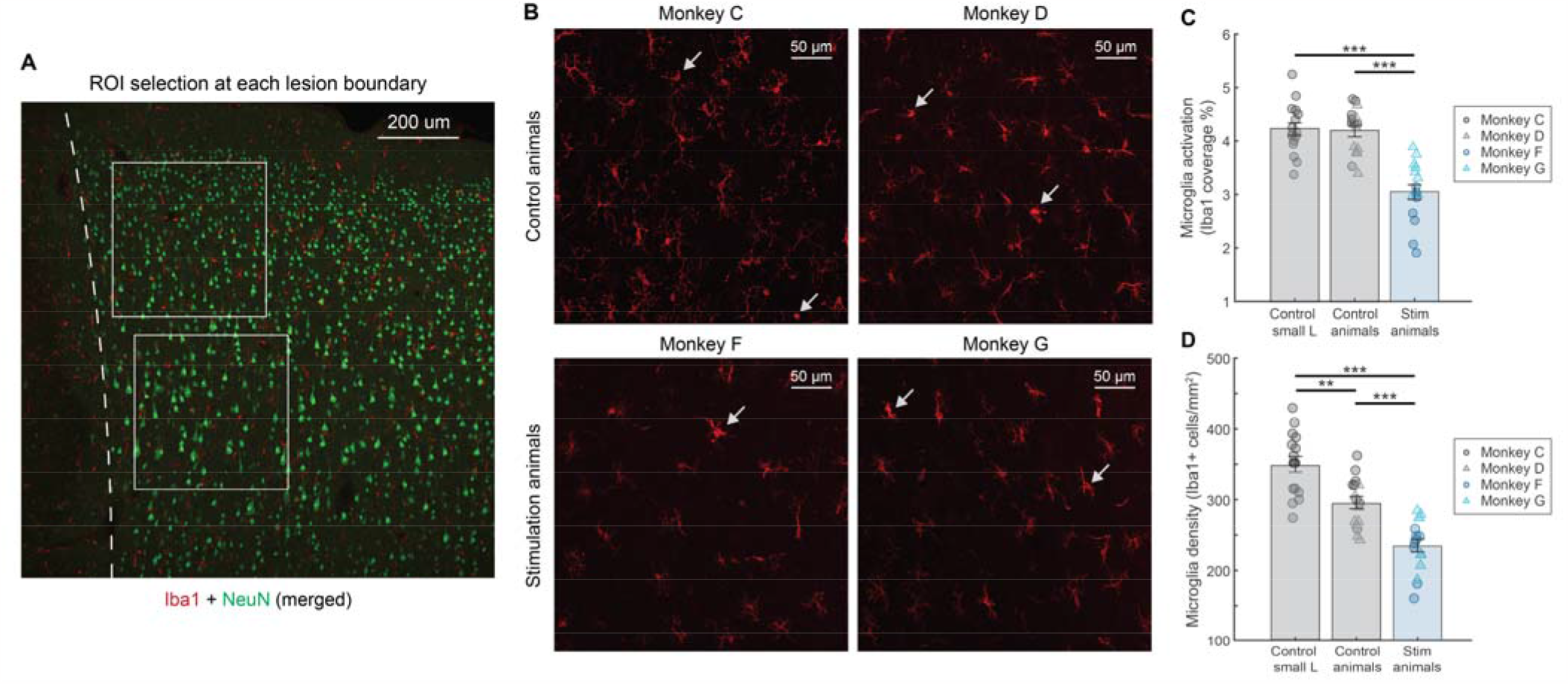
Immunohistochemistry staining of Iba1 protein (microglia marker). **A**. Confocal image of an ipsilesional coronal section stained with Iba1 (red) and NeuN (green). Dashed line represents the lesion boundary. White squares represent two sample ROIs selected for microglial analysis. **B**. Sample ROIs of Iba1 staining in each monkey at the lesion boundary. White arrows point to examples of morphologically activated microglia. **C**. Summary of microglia activation as measured by percentage coverage of Iba1-positive pixels for the control (monkeys C and D) and stimulation group (monkeys F and G) at the lesion boundary. Two smaller lesions in monkey C are used as additional controls. One-way ANOVA detected a significant difference in microglia activation among the three groups (p<0.001). Post-hoc Bonferroni test suggested that stimulated animals have significantly lower microglia activation comparing to both controls (***: p<0.001). **D**. Summary of Iba1-positive microglia density for the control and stimulated group at the lesion boundary. One-way ANOVA detected a significant difference in microglia density among the three groups (p<0.001). Post-hoc Bonferroni test revealed that stimulated animals have significantly lower microglia density comparing to both controls, while peri-infarct regions at smaller lesions have significantly higher microglia density (***: p<0.001, **: p<0.01).

## Discussion

In this study, we revealed possible mechanisms of stimulation-induced neuroprotective effects after acute ischemic stroke by combining the latest technology in electrophysiology and histology with a unique NHP stroke model. Our results indicate that early electrical stimulation decreases the extent of neuronal cell death by reducing peri-infarct depolarization, excitotoxicity, and inflammation in the sensorimotor cortex of NHPs. These findings suggest that using electrical stimulation to protect the brain and reduce tissue damage during acute ischemic stroke could play a critical role in alleviating the global burden of stroke, as infarct size plays a critical role in determining not only patient mortality, but also the functional outcome of chronic stroke rehabilitation strategies.

We utilized the photothrombotic method to produce focal ischemic lesions in NHPs. In comparison to other interventions for generating infarcts (42), our method is less surgically challenging and allows for more reliable control of the location and size of infarcts across animals by implementing the same aperture, intensity, and duration of light illumination as demonstrated in our previous work (31). However, it should be noted that photothrombotic infarcts do not model all aspects of acute stroke in humans. For example, photothrombotic lesions only generate very narrow penumbras and are largely limited to the superficial layers of the cortex, thus not covering other stroke pathways such as those dominated by subcortical and white matter damage (43). Nevertheless, we were able to generate controlled focal lesions in the sensorimotor region of NHPs using this method, while simultaneously collecting ECoG recordings from the impacted brain to monitor neural activity changes. For the lesion volume estimation and electrophysiology analysis, we compared the results from control monkeys D and E to the stimulated monkeys F and G, which were lesioned using identical illumination parameters that have shown to induce predictable infarct sizes (31). However for monkey E, we observed very poor immunofluorescence signals while having normal density of Nissl-stained cells over the exposed sensorimotor cortex. It is unclear whether this lack of immunoreactivity was caused by surgical procedures, perfusion fixation, or subsequent tissue handling. Therefore, we added monkey C as an additional control and excluded monkey E for all immunohistochemistry procedures. Monkey C received light illumination through various apertures during lesion induction. As a result, it not only serves as a no-stimulation control for the immunohistochemistry analysis, but also contains multiple smaller lesions to help distinguish the cellular effect of electrical stimulation from the direct impact of variability in photothrombotic infarct sizes.

We applied repeated electrical stimulation adjacent to the lesion on the ipsilesional cortex, 60 minutes after lesion induction in monkeys F and G. The stimulation train contains 5 Hz bursts of biphasic pulses, similar to the theta burst stimulation (TBS) pattern of transcranial magnetic stimulation (TMS) protocols that are widely adopted in the clinic (44). In past studies, continuous TBS protocols have been reported to have an inhibitory effect on synaptic transmission and cortical excitability in human subjects (45, 46). However, one major difference is that our stimulation paradigm used five pulses at 1 kHz in each burst in contrast to the three 50-100 Hz pulses used in traditional TBS protocol. Stimulation via high frequency pulses at greater than ∼200 Hz have been shown to have an inhibitory effect on neuronal firing rates (47). Meanwhile, similar patterns of pulse train have also been used for paired-pulse conditioning through intracortical microstimulation and were shown to promote Hebbian-like plasticity in rodents (48). Given that our ECoG recordings showed decreasing gamma power over the course of stimulation but not in control animals, and that gamma activity in ECoG has been shown to correlate with neuronal firing (34), it is possible that applying theta bursts of electrical stimulation over the sensorimotor cortex decreased neuronal activation and network excitability through mechanisms similar to those seen in continuous TBS and high frequency pulse trains which induce hyperpolarization and synaptic depression (45, 49). As smaller infarct volume was observed in the stimulated animals along with the downregulated neural activity, our results provide evidence that cortical stimulation prevents tissue damage by reducing excessive depolarization and glutamate-mediated excitotoxity adjacent to the lesion during the ischemic cascade. These results are also encouraging because the hypothesized neuroprotective mechanism is supported by the therapeutic strategies of many pharmacological agents designed for acute ischemic stroke, which aims to attenuate excitotoxicity and restore the balance between oxygen supply and energy consumption by inhibiting neuronal excitability (5, 6). In addition, adopting electrical stimulation as a therapy for acute stroke can be more advantageous for patients suffering severe side-effects or allergy due to the neuroprotective drugs.

We performed immunohistochemistry staining against c-Fos and Iba1 to assess neuronal activation and microglial response in the perilesional tissues of the control and stimulated animals respectively. Since microglial response after acute stroke is reflected by both cell migration to the peri-infarct region and morphological activation through cell body enlargement and process thickening (50), we used two measures to quantify the inflammatory response: Iba1-positive cell density and percentage area coverage by Iba1-positive pixels. Between these two measures, percentage area coverages offers a more accurate estimation of the level of inflammation since it accounts for both microglia accumulation and activation within a given region. Importantly, the reduced c-Fos activity and microglial response around the lesion boundary offered us complimentary information when combined with the electrophysiology results above, suggesting that electrical stimulation reduced both the level of cortical depolarization and neuroinflammation for tissues undergoing the acute ischemic cascade. These findings confirmed what has been reported before for rodents receiving cortical stimulation via bipolar electrodes (27, 30) or cathodal transcranial direct current stimulation (C-tDCS) (25, 26), in which stimulation decreased tissue damage by inhibiting apoptosis, neuroinflammation, and peri-infarct depolarization during acute stroke. In addition, the decrease in c-Fos immunoreactivity in the stimulated monkeys suggests that the downregulation of ECoG activity is not a manifestation of the harmful cortical spreading depression, since an upregulation of c-Fos has been shown to correlate with sustained depolarizations and the subsequent spreading depression in the presence of focal ischemia (51). Combined with the reduction in both lesion size and microglial accumulation, our findings suggest that electrical stimulation applied one hour after stroke onset offered inhibitory and protective effects instead of exacerbating tissue damage attributed to spreading depolarization as previously described for early sensory stimulation (23), making this stimulation protocol a safe and promising treatment option for acute ischemic stroke.

In the future, additional experiments can be performed to validate the reduction in ischemic infarct volumes statistically between control and stimulation groups. In addition, applying the same lesioning toolbox and acute electrical stimulation to monkeys chronically implanted with our multi-modal interface can allow us to study the impact of stimulation well beyond the acute window. Combined with the sensorimotor behavioral capabilities of NHPs, these experiments will provide us with valuable insights into the mechanisms of stimulation-induced neuroprotection and its effect on functional recovery from days to months after stroke onset. Investigating the chronic effect of localized electrical stimulation in a translatable NHP stroke model with consistent lesion size and location can help bridge the gaps between rodent and human studies that hindered the success of past clinical trials on neuroprotective therapy (9, 10). Furthermore, this stimulation paradigm can also be explored in conjunction with reperfusion and other pharmacological therapies to further improve the neurological and functional outcome after ischemic stroke in the clinic.

## Materials and Methods

### Animals subjects and surgical procedures

All animal procedures were approved by the University of Washington Institutional Animal Care and Use Committee and were in accordance with the National Research Council’s Guide for the Care and Use of Laboratory Animals. All our animals were housed and maintained at the Washington National Primate Research Center (WaNPRC) accredited by the American Association for Assessment of Laboratory Care (AAALAC). This study used five adult macaques (Control group: monkey C - *Macaca mulatta*, 10.3 kg, 16 years, female; monkey D - *Macaca nemestrina*, 12.8 kg, 14 years, female; monkey E - *Macaca nemestrina*, 13.10 kg, 14 years, female; Stimulation group: monkey F - *Macaca nemestrina*, 13.8 kg, 14 years, female; monkey G - *Macaca nemestrina*, 14.6 kg, 7 years, male) that were part of the Tissue Distribution Program (TDP) at the WaNPRC, which aims to conserve and fully utilize the NHPs no longer needed for other experiments.

### Surgical Procedures and Induction of Focal Ischemic Lesions

Using standard aseptic technique, the five macaques (Monkey C, D, E, F, G) were anesthetized with isoflurane and placed in a stereotaxic frame. The animals’ temperature, oxygen saturation, heart rate, and electrocardiographic responses were monitored throughout the procedure. Bilateral craniotomies and durotomies (25 mm diameter) were performed using stereotaxic coordinates that target the sensorimotor cortices (52). To protect the exposed brain, a transparent silicone artificial dura (monkey C) or a semi-transparent multi-modal artificial dura (monkey D to G) fabricated using previously described methods (53, 54) was implanted bilaterally on top of the sensorimotor cortex, offering optical and electrical access during subsequent procedures. For animals in both the control (monkeys C, D, E) and stimulation groups (monkeys F, G), we induced ischemic lesions using the photothrombotic technique (31, 33), which produces focal infarcts by photo-activation of a light-sensitive dye (Rose Bengal). Upon illumination, the intravenously administered dye produced singlet oxygen that damaged endothelial cell membranes, causing platelet aggregation and interrupting local blood flow. To control the infarct size and location, we placed an opaque silicone mask on top of the artificial dura. In monkey C, the mask contains circular apertures of different diameters (0.5, 1.0, and 2.0 mm) to induce control lesions in various sizes. In monkey D to G, the mask has a single aperture located in the center (diameter 1.5 mm) and was placed on one hemisphere (monkeys D, F, and G: left hemisphere; monkey E: right hemisphere). After the mask was in place, each animal was injected with Rose Bengal (20mg/kg) for 5 minutes as we started illuminating the ipsilesional cranial window for 30 minutes through the aperture using an uncollimated white light source.

### Electrophysiology recording and electrical stimulation

All electrophysiology recordings and electrical stimulation were performed with Grapevine Nomad processors, four Nano front ends (Ripple Neuro, Salt Lake City, UT), and our customized large-scale multi-modal interface. The design and characterization of this interface with 32 embedded ECoG electrodes can be found in our previous work (52, 53). A skull screw close to the midline and anterior to the ipsilesional cranial window was used as an electrical ground for subsequent recordings. All animals were transitioned from isoflurane to urethane anesthesia prior to ECoG recordings and stayed on urethane until the end of the experiment to allow reliable monitoring of neural activity. In control monkeys D and E, we collected ECoG data bilaterally at 30 kHz sampling frequency, including 30 minutes of baseline before photothrombotic lesioning, 30 minutes during illumination, and up to 3 hours post lesioning to monitor the extent of neuronal damage and network dynamics around the injury site. In stimulated monkeys F and G, we followed the same recording timeline for baseline and illumination periods, and recorded spontaneous neural activity for 1 hour after lesioning. We then applied electrical stimulation through a single ECoG electrode approximately 8 mm medial to the lesion center on the ipsilesional (left) hemisphere. We delivered the stimulation trains in 6 blocks lasting 10 minutes each, with 2-minute recordings of spontaneous activity in between the blocks to track changes in neurophysiology as stimulation continued. The stimulation trains had a 5 Hz burst frequency and 5 biphasic charge-balanced pulses at 1 kHz within each burst. The stimulation amplitude was 60 μA and pulse width was 200 μs per phase with 50 μs inter-phase interval.

### Electrophysiology Data Analysis

Signal power calculation of both lesion and non-lesion electrodes was conducted in MATLAB (MATLAB R2022b, MathWorks). After down-sampling the signal from 30 kHz to 1 kHz, the signals were notch filtered at 60, 120, 180, and 240 Hz. Channels with a power spectral density that did not exhibit the expected 1/f ll curve were excluded from further analysis. As a result, one channel was excluded for monkeys D and G, two channels for monkey E, and fourteen channels for monkey F. Next, we filtered the signal into distinct frequency bands including gamma (30-59 Hz) and high gamma (60-150 Hz). The signals were then split into two-minute blocks separated by ten minutes across the pre-stroke baseline, post-stroke, and post-stimulation recording periods. We then identified signal artifacts by normalizing each signal and excluded the time points in the non-normalized signal for which the normalized signal amplitude exceeded 25 standard deviations from the mean of the signal. Prior to calculating signal power, we normalized the signal of every two-minute window within each channel by dividing by the maximum of the absolute value of the baseline signal. We then calculated the average signal power of each channel for each two-minute window by squaring the normalized signal and dividing by the elapsed time. To compare power across each two-minute time window, we conducted paired sample t-tests with Bonferroni corrections for multiple comparisons (family-wise error rate of 0.05). Pairwise comparisons of power distributions were only made between baseline and each subsequent two-minute window and between 50 min post-stroke and each subsequent two-minute window following stimulation.

### Histology, Lesion Reconstruction, and Size Estimation

At around 4 hours after the stroke was induced, animals were deeply sedated and transcardially perfused with phosphate buffered saline (PBS) followed by 4% paraformaldehyde (PFA). The brains were harvested and post-fixed by immersion in 4% PFA for 24 to 48 hours. A coronal block containing the lesioned region was dissected using a custom matrix and then stored at 4 °C in 30% sucrose in PBS. To prepare for staining, the block was frozen and sectioned into 50 μm thick coronal sections using a sliding microtome (Leica). Sliced sections were stored at 4 °C in PBS with 0.02% sodium azide.

To evaluate the extent of ischemic damage and neuronal death, we mounted a rostrocaudal series of coronal sections with approximately 0.45 mm separation and then performed Nissl staining on the mounted sections using Thionin acetate. Nissl-stained sections were scanned and registered using a custom software in MATLAB (2019b, MathWorks) for alignment and three-dimensional reconstruction. The registered images were smoothed, binarized, and went through edge detection so that lesion boundaries on each slice can be identified and visualized within the reconstructed cortex. A more detailed description of the lesion edge detection and reconstruction method can be found in our previous publication (31). The widths and depths of each lesion within the representative coronal sections were also calculated based on the detected boundary and image resolution. Linear interpolation was used between sections to estimate the lesion volume in each animal. Furthermore, to combine histological findings with electrophysiology recordings, we overlayed the surgical image taken for each animal on top of its reconstructed cortex based on the location of sulci and other distinct anatomical features. The overlayed images allowed us to register the ECoG electrodes that fell within the estimated lesion area, and classify them into lesion and non-lesion groups respectively for the subsequent electrophysiology analysis.

### Immunohistochemistry

To evaluate the neuronal activation and neuroinflammatory response near the ischemic lesion, we performed immunostaining with antibodies against c-Fos protein and the microglia/macrophage-specific calcium-binding protein Iba1 respectively. Coronal sections around the lesion in monkeys C, D, F, and G (4 sections around each lesion) were co-stained with either neuronal nuclear protein NeuN and c-Fos or NeuN and Iba1. Monkey E was excluded for all immunohistochemistry analysis due to lack of immunoreactivity at the cranial window. To replace monkey E, we stained and analyzed tissues from monkey C as an additional control. In monkey C, we only selected coronal sections with a single lesion either comparable to or smaller than those in monkeys D to G for subsequent analysis. To prepare tissues for staining, coronal sections were first rinsed in 1x PBS and incubated in 1% NaBH_4_ solution for 1 hour to reduce background autofluorescence, after which they were washed with PBS and incubated in normal donkey serum (NDS) blocking solution (10% NDS and 0.1% triton-X100 in PBS) overnight at 4°C. Sections were then incubated in primary antibodies including either rabbit anti-c-Fos (Abcam ab190289, 1:500 dilution) or goat anti-Iba1 (Abcam ab5076, 1:1000 dilution) plus mouse anti-NeuN (Millipore Sigma MAB377, 1:500 dilution) in NDS blocking solution at 4°C for ∼72 hours. Sections were then rinsed in PBS and incubated in secondary antibody mixtures containing either donkey anti-rabbit antibody (Invitrogen #A10042, 1:500 dilution) or donkey anti-goat antibody (Invitrogen #A-11057, 1:500 dilution) conjugated with Alexa Fluor 568, plus donkey anti-mouse antibody conjugated with Alexa Fluor 488 (Invitrogen # A-21202, 1:500 dilution) and DAPI at 4°C overnight. This was followed by rinsing sections in PBS for 5 times and then in 1:1 Glycerol-PBS solution for 10 minutes. Sections were mounted onto slides using DABCO mounting media and later imaged using a Leica widefield fluorescence microscope or a Nikon A1R HD25 laser scanning confocal microscope.

### c-Fos and microglia activation analysis

To estimate the density of c-Fos positive cells, we took four images per coronal section at both sides of the lesion boundary identified from NeuN expression using a 10x objective through the widefield fluorescence microscope. We then used custom algorithms in MATLAB to perform binarization with adaptive thresholding followed by image segmentation. The segmented cells with positive c-Fos immunoreactivity outside of the lesion were counted automatically and the cell density was computed based on the size of counted area in each image. To evaluate the level of microglial reactivity, we imaged both sides of the lesion boundary using a 20x objective through the laser scanning confocal microscope. For each region of interest (ROI) within 1 mm from the lesion boundary, we took z-stack images with 20 μm thickness and created a maximum intensity projection per ROI. The projections were converted to 8-bit and inverted so that dark pixels represent Iba1 signals. We then used ImageJ to denoise each ROI by subtracting the background and removing pixel intensity outliers. The ROIs were converted to binary by adjusting the threshold to one standard deviation below the average pixel intensity and filtering out objects below 150 pixels in size. The percentage area covered by dark pixels, a widely used Iba1 reactivity measure (55), was then calculated to estimate the microglia activation level. Finally to quantify microglial accumulation, we used the same binarized images and counted the number of segmented cells in each ROI to estimate the average microglia density at the lesion boundary for each animal.

## Acknowledgment

We thank Toni Haun, Devon J. Griggs, Christopher English, Britni Curtis, and Sandi Thelen for their help with animal care, experimental preparation, and assistance during surgeries. We also thank Aryaman Gala and Mona Rahimi for their help with histological procedures and preliminary analysis. This study is supported by the National Institute of Neurological Disorders and Stroke of the National Institute of Health (NIH 1R01NS119395-01, A.Y. and J.Z.), the Washington National Primate Research Center (WaNPRC, P51 OD010425), the American Heart Association, the National Science Foundation Graduate Research Fellowships Program (K.K.), the Washington Research Foundation, and the Weill Neurohub (J.Z.).

